# Hierarchical Predictive Processing during Natural Reading

**DOI:** 10.64898/2026.06.24.734402

**Authors:** Lucas Y. H. Chan, Olaf Dimigen, Benjamin Gagl, Urs Maurer

## Abstract

Key neural mechanisms underlying reading, especially in the context of sentences and texts, remain elusive. An influential theory states that the brain rapidly generates a hierarchy of probabilistic predictions about upcoming words at multiple levels of linguistic representation. When encountering a word, a reader integrates sensory input with the linguistic predictions, and rely on the resulting prediction error signals to achieve efficient comprehension. Here, we tested this hierarchical predictive account with a natural reading task while EEG and eye movements were simultaneously recorded. We find distinct prediction error signals attributable to word contextual, lexical, and orthographic levels of predictions. These prediction error signals exhibit a temporal order. Our results also indicate that context-based predictions rapidly constrain orthographic-level predictions. We thus provide evidence for a hierarchical predictive process central to efficient natural reading.

---

We comprehend meaningful messages by rapidly scanning words, sentences, and texts. This ability to read is a foundation to human communications that traverse distances in time and space. While skilled readers can read with remarkable efficiency (*1, 2*), the underlying neural mechanism remains elusive. The Hierarchical Predictive Processing hypothesis (HPP, *3–5)* is a highly influential idea for understanding neural mechanisms, including those underlying language comprehension (*6–8*). HPP posits that the brain generates top-down probabilistic predictions about upcoming sensory information on multiple processing levels. Prediction error signals are in turn elicited at each level that scale with the magnitude of difference between the prediction and sensory input encountered. Prediction error, which represents sensory information not predicted by the brain, enables neuronal processing of sensory input with high efficiency (*4, 9*).

For language comprehension, recent studies have shown evidence of HPP on high-level (e.g., context) and low-level (e.g., visual and auditory features) linguistic representations respectively (*10–14*). Furthermore, HPP also suggests the integration between high and low level representations as a core mechanism of comprehension (see *6*). This is supported by studies which showed an optimizing effect of contextual prediction on low-level sensory inputs (*15–17*). However, much current evidence is established from listening comprehension (*10, 11, 14,17*) or visual single-word recognition tasks (*12, 13, 15, 16*). It remains unknown whether the comprehension of written text also engages a similar process, particularly in natural reading conditions.

For decades, the neural mechanisms underlying sentence reading have been mainly studied in highly simplified laboratory tasks that involve the passive visual presentation of sentences one word at time (rapid serial visual presentation, RSVP). While the RSVP paradigm avoids certain measurement artifacts, it is very much different from typical reading conditions (see *18*). For example, natural reading requires the reader to actively move their eyes through the text and allow the reader to perceive visual information in foveal and parafoveal regions. An alternative method, which allow the study of brain activations in more natural conditions, is the simultaneous recording of electroencephalography (EEG) and eye movement. This method, combined with recent advances in artifact correction (*19*) and statistical modelling (*20*), enables the synchronization between EEG and eye movement data with sub-millisecond precision, and allows us to reliably track the neural responses aligned to eye fixation onsets during natural reading (fixation-related potentials, FRPs, *21*).

Here, we report a study in which subjects participated in a natural reading task while their EEG and eye movement were simultaneously recorded with high temporal resolution (Fig. 1A and 1B). We set out to investigate HPP at three levels: contextual, lexical, and orthographic word representations. If natural reading involves hierarchical predictions, we expect fixation-related brain activity to show: (1) distinct prediction error signals at each level of the hierarchy, (2) a temporal order between the prediction error signals, and (3) that high-level predictions dynamically constrain low-level predictions.

**Fig. 1.**
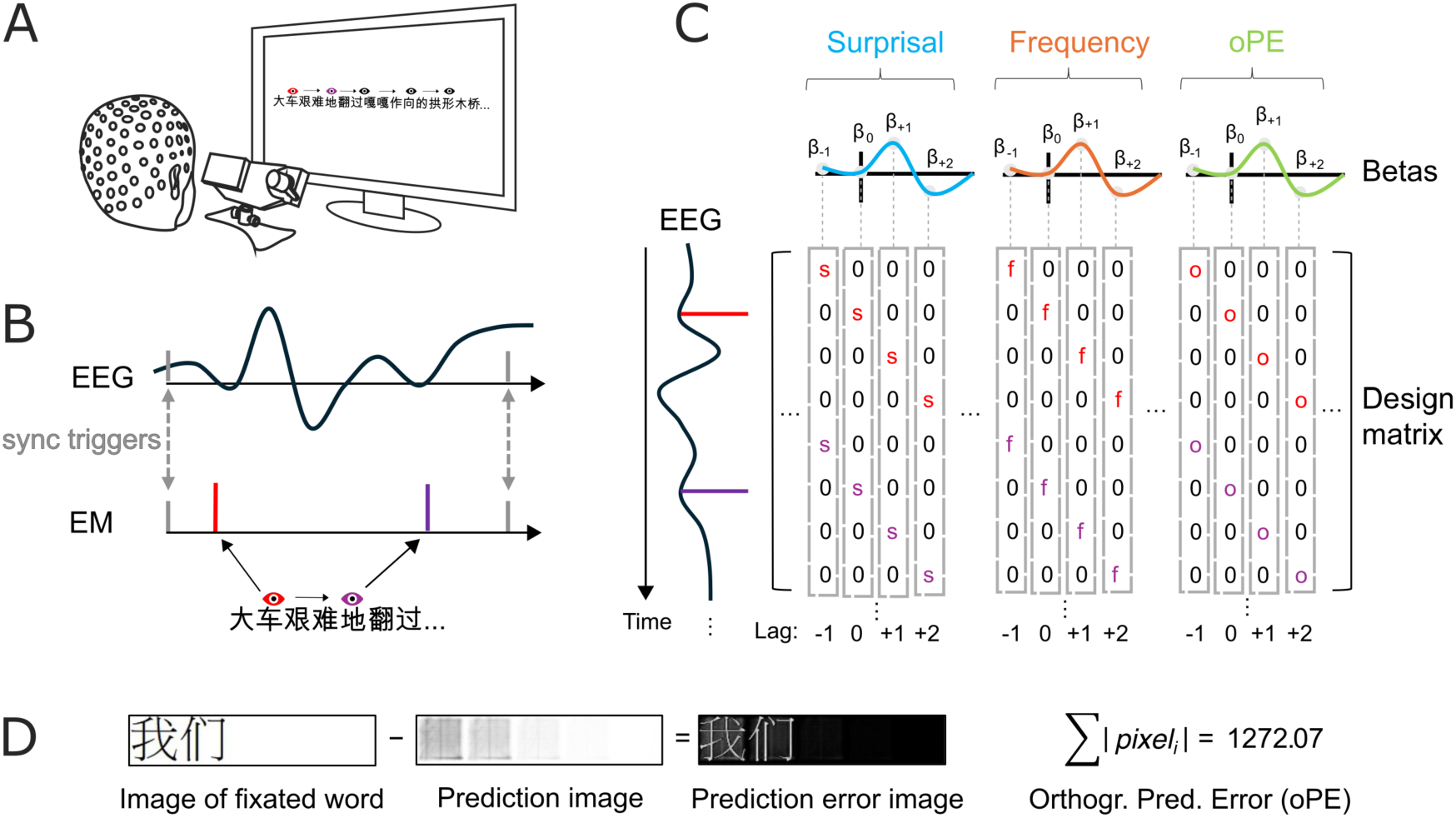
The natural reading task, regression-based fixation-related potentials (rFRPs), and Orthographic Prediction Error (oPE). **(A)** During the experimental task, participant read novel excerpts while their EEG and eye-movements were simultaneously recorded. **(B)** The EEG and eye-movements data were synchronized offline based on common triggers between the two datasets. **(C)** The fixated word’s surprisal, frequency, and oPE (along with other control variables) were used to predict the EEG data in a regression model. The design matrix was time-expanded so that each of these predictors modelled from 200 ms before and 800 ms after the fixation events. The resulting beta coefficients from each predictor (rFRP) then entered into further analysis. **(D)** oPE is the pixel-by-pixel difference between the prediction image and the image of the fixated word (*13*). The prediction image was derived by averaging 23,854 word images, thus depicting the "average" word orthographic pattern of the language. When computing oPE, the image of the fixated word was subtracted from the prediction image which resulted in a prediction error image. oPE is the sum of all absolute pixel intensities (on a continuous scale from 0 to 1 where 0 = Black and 1 = White) of the prediction error image.

Prediction error signals at each level were modelled (see Fig. 1C) using word *surprisal*, *lexical frequency*, and the *orthographic prediction error* (oPE, *13*). A word’s surprisal value represents the error signal based on all information provided by the text’s prior context (such as semantics, syntax, morphological expectations). Here, it was defined as the negative-log context-dependent probability estimated by the Chinese-Llama 2 Large Language Model (LLM, *22*). Contextual probabilities derived from LLM have shown to be comparable to those estimated from the classic cloze procedure (*23*). Next, lexical frequency measures a word’s probability of occurrence independent of the prior context (e.g., low-frequency words are less likely to occur in a language). The inverse of lexical frequency represents the error signal based on general statistical regularities of words contained in the language and also in the participant’s mental lexicon. Lexical frequency was retrieved from the SUBTLEX-CH corpus (*24*). Finally, oPE (*13*) is the pixel-by-pixel difference between an orthographic prediction image and the actual orthographic input (Fig. 1D). The orthographic prediction image was derived by averaging pixel intensities across 23,854 word images. The summed oPE value represents the error signals elicited by word predictions based on the participant’s knowledge about the language’s orthographic patterns.

To this end, we hypothesized that on each of the three levels, any mismatch between the reader’s prediction and the actual word input will elicit a prediction error signal that scales with the degree of the deviation. EEG should detect these signals on the scalp and exhibit distinct temporal and spatial patterns for prediction error signals arising at each level.

## Results

### Distinct prediction error signals at three levels of word representation

We analyzed natural reading EEG data using the regression-ERP framework (*25, 26*) with the *unfold* toolbox in MATLAB (*20*). Importantly, this so-called linear deconvolution framework allows us to disentangle and isolate the influences of different word properties and from successive eye fixations on the EEG during natural reading (*27*). Within the model, for each fixation on a word, we used the fixated word’s surprisal, frequency, and oPE to model the EEG response from 200 ms before to 800 ms after fixation onset. To control for potential confounds, lexical and oculomotor covariates, known to influence the fixation-related EEG, were included as regressors in the model. Specifically, it controlled for incoming saccade amplitude, the fixated word’s length, and the brain response elicited by the initial onset of the sentence stimulus. Correlations between the regressors are available in the supplementary materials (Table S1). The beta coefficients (i.e., regression-based FRP, rFRP) of each of the three predictors of interest were then entered into further analyses.

#### Surprisal

We first tested whether our model captured prediction error signals at the level of contextual word prediction. Indeed, cluster-based permutation test on the surprisal rFRP revealed a significant effect (peak significance at 237 ms, *p* = .004). To inspect the time course of the effect, we computed Global Field Power (GFP) and Global Map Dissimilarity (GMD, *28*) of the predictor’s rFRP. GFP and GMD measure, respectively, the grand-average topographic map strength and stability across time. Results showed a relatively strong and stable topographic distribution approximately from 155 ms to 552 ms (Fig. 2A). Within this period, topographic maps showed that higher word surprisal elicited increased negativity first at left-lateralized occipital regions and then at centroparietal regions, as well as increased positivity at left-frontal regions throughout the entire period. The topographic distribution and the rFRP at centroparietal electrodes (Fig. 2B) resemble the N400 effect (see *29*) found in response to reading words with low cloze probability (e.g., *30–32*) or high surprisal (e.g., *33, 34*). The slightly earlier peak in the rFRP, compared to ERP studies, is in accordance with the N400 effect found in natural reading studies (e.g., *21, 35*). The increased positivity at left-frontal regions is also consistent with results reported in some studies (e.g., *30, 32, 36*), potentially due to the use of average instead of mastoid EEG reference.

**Fig. 2.**
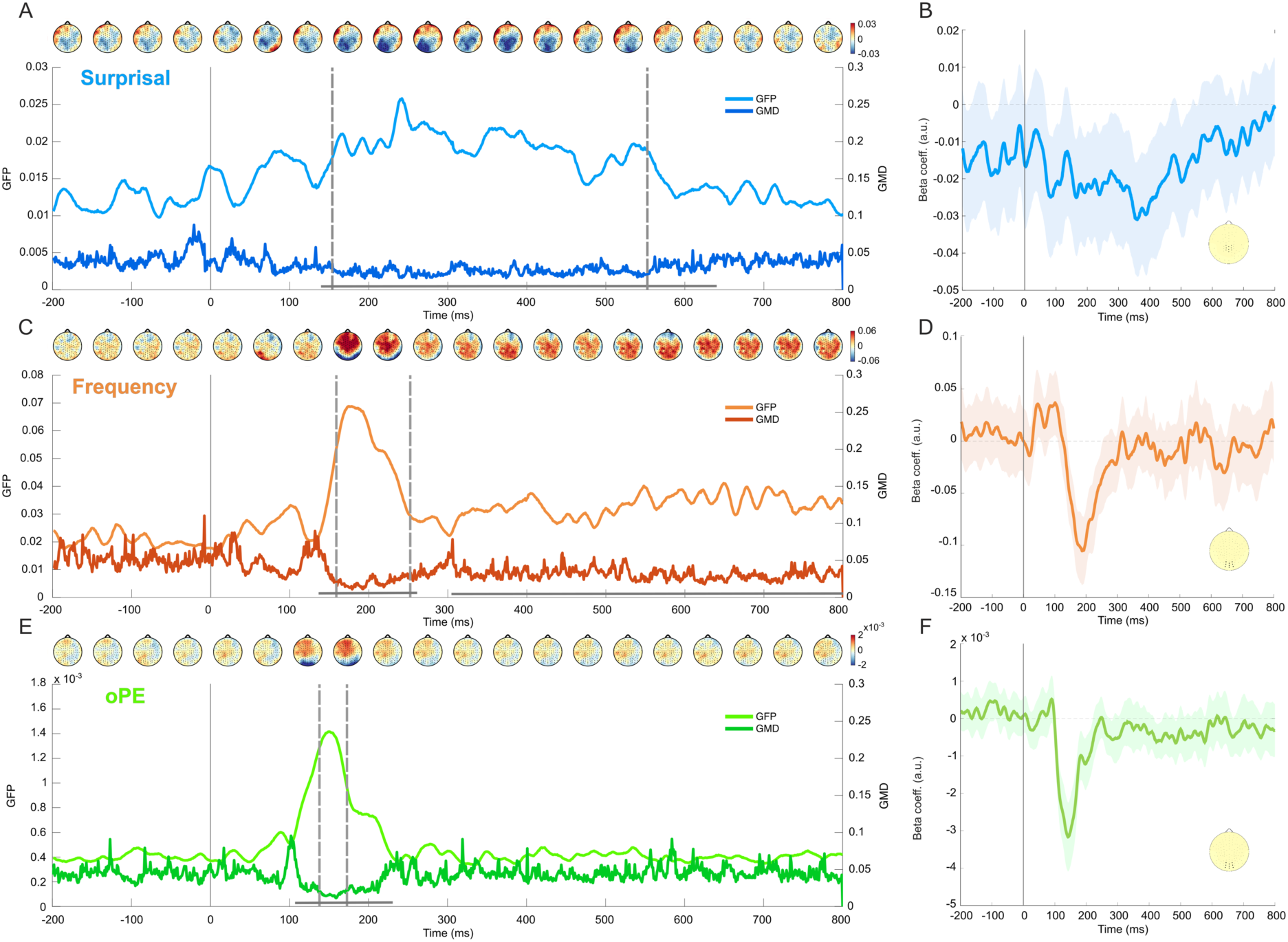
Neural correlates of three levels of prediction error signals in fixation-related brain potential. The left column shows the grand-average Global Field Power (GFP), Global Map Dissimilarity (GMD), and topographic maps of the **(A)** surprisal, **(C)** frequency, and **(E)** oPE effects as a function of time. GFP and GMD respectively indicate the strength and stability of each effect across time. The dotted lines mark an approximate period of high GFP and low GMD which shows a period of strong and stable effect. Grey horizontal lines indicate significant clusters with *p* < .05. The right column shows the grand-average rFRP of the **(B)** surprisal effect averaged across midline parietal electrodes, **(D)** frequency effect averaged across midline occipital electrodes, and **(F)** oPE effect averaged across midline occipital electrodes. They respectively show effects at the N400, N250, and N1 components found in traditional ERP studies. Shaded areas represent 95% confidence interval.

#### Frequency

Next, we sought to see whether predictions are also made based on word lexical frequency. We found significant clusters in the rFRP of the word frequency predictor (peak significance at 189 ms, *p* < .001). A high GFP and low GMD of the frequency effect was observed approximately from 160 ms to 250 ms, indicating a reliable effect in this period (Fig. 2C). Topographic maps within this period showed that less frequent words elicited increased negativity at occipital regions, and increased positivity throughout central and frontal regions. These results suggested that lower word frequency is related to more negative (or less positive) amplitude near the N1 component (Fig. 2D). This is consistent with ERP studies that showed low frequency words elicited a more negative N1/N170 (e.g., *37, 38*), or a less positive P2 (e.g., *36*), when compared to high frequency words. Similar effects were also found in FRPs (*39–42*). On the other hand, some studies also reported the N400 effect elicited by low frequency words (e.g., *42, 43*). However, we did not find reliable evidence that suggested an effect within this time window. One likely reason is that our surprisal predictor had captured most of the N400 effect.

#### Orthographic Prediction Error

Lastly, we found significant clusters in the rFRP of the oPE predictor (peak significance at 142 ms, *p* < .001). There were high GFP and low GMD approximately from 138 ms to 171 ms (Fig. 2E). Within this time window, topographic maps showed that higher oPE elicited increased negativity at posterior-occipital regions (Fig. 2F), and increased positivity at centroparietal and frontal regions. The topographic distribution resembled that of previously reported oPE effects, in which higher oPE elicited stronger negativity at occipital areas (*13*). Our result is also largely consistent with previous research suggesting that the N1/N170 component reflects early orthographic processing (e.g., *44–46*).

P-value maps of the three predictors are available in supplementary materials (Fig. S1). As control analyses, we also tested regression models with the oPE predictor normalized for word length and models that were restricted to first-pass fixations on one-character words. The results of the control analyses were comparable to those reported above, indicating that the effects were not influenced by factors such as re-fixations or the collinearity between oPE and word length. Results of the control analyses (Fig. S2) and analyses on gaze durations (Table S2) are available in supplementary materials.

### A temporal order of prediction error signals among the three levels

We showed in the previous section that the brain generates distinct prediction error signals at different levels of linguistic representation. Next, we sought to confirm whether there is a temporal order among these signals. We computed two measures of jackknife-based effect latencies (*47*). First, for each effect, we defined the peak effect latency as the time at which the correlation was maximal between the topographic map at grand-average GFP maxima (Fig. 3A) and jackknife-based rFRP maps from 0 ms to 400 ms (*48, 49*). Second, we defined the "central" effect latency as the centroid of each jackknife-based GFP waveform within the same time window. We tested the latency differences with linear mixed models and found significant differences in both peak and central effect latencies among the three effects (both *p* < .001, Fig. 3B). For peak effect latencies, pairwise comparisons revealed that the peak effect of oPE (*M =* 149.95, *SD* = 0.50) occurred significantly earlier than the peak effect of frequency (*M =* 175.03, *SD* = 0.53), *t*(117) = 234.0, *p* < .001, and surprisal (*M =* 242.0, *SD* = 0.39), *t*(117) = 859.0, *p* < .001. The peak effect of frequency was also significantly earlier than that of surprisal, *t*(117) = 625.0, *p* < .001. With regard to central effect latencies, we also found that the oPE effect (*M =* 185.15. *SD* = 0.90) preceded that of frequency (*M =* 207.41. *SD* = 1.00), *t*(78) = 93.6, *p* < .001, and surprisal (*M =* 213.54. *SD* = 1.38), *t*(78) = 119.4, *p* < .001. Finally, the central effect of frequency occurred earlier than that of surprisal, *t*(78) = 25.8, *p* < .001. These results suggest a temporal ordering among the three prediction error signals, from orthographic to lexical, and finally, to contextual word predictions.

**Fig. 3.**
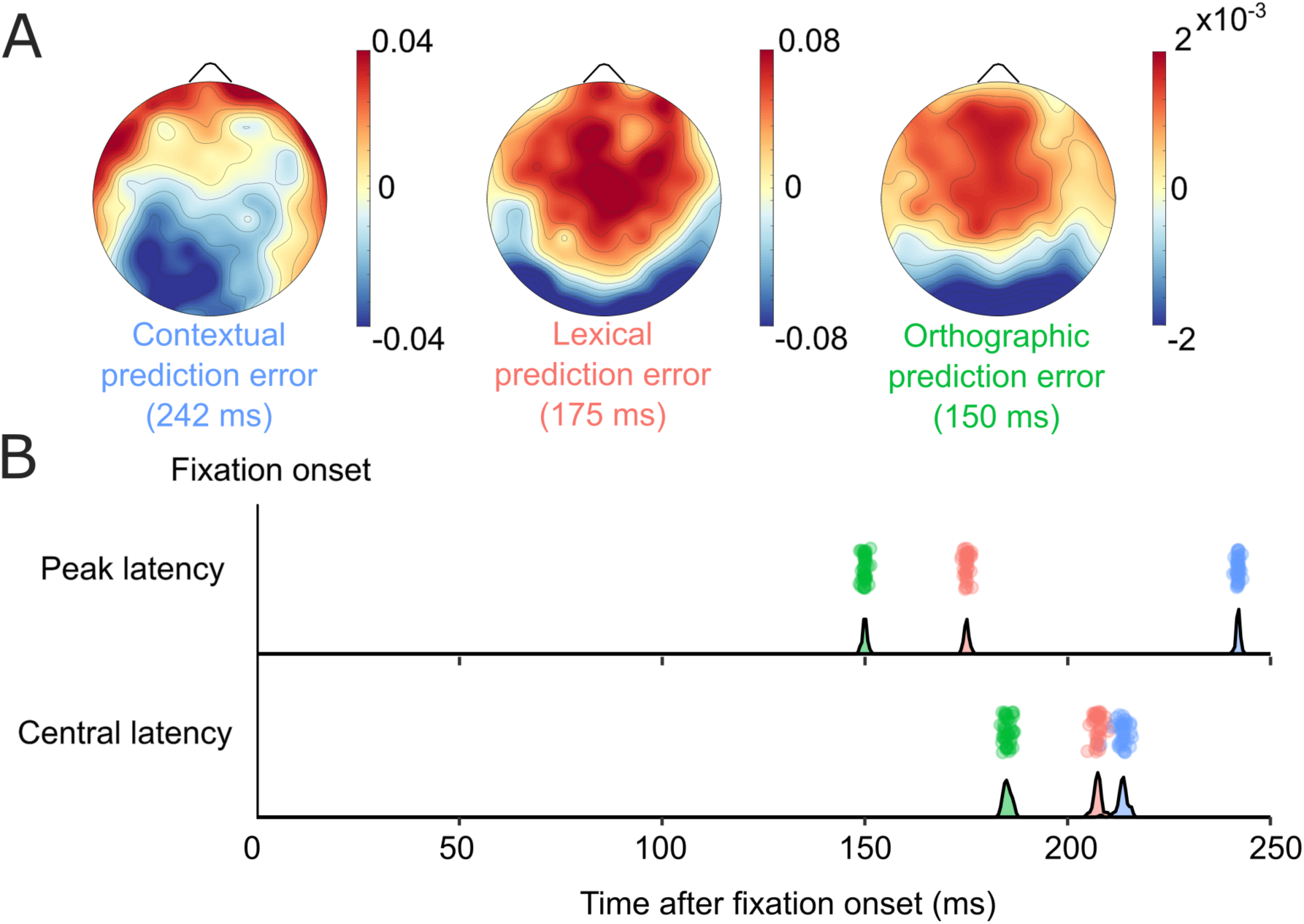
Topographic maps of the three effects at their respective GFP peak, and their peak and central effect latencies after fixation onset. **(A)** Topographic maps taken at 242 ms of the surprisal effect, at 175 ms of the frequency effect, and at 150 ms of the oPE effect. These maps were used as template maps for identifying peak latencies. **(B)** The peak latencies (upper row) and central latencies (lower row) among the three effects. Each dot represents the latency value of a jackknife-based waveform. The density curves on the x-axes show the distribution of latencies among each effect’s jackknife-based waveforms. Both peak and central latencies show a temporal order among the three prediction error signals: from oPE, to lexical prediction error, and finally contextual prediction error. Green: oPE effect latencies; Orange: frequency effect latencies; Blue: surprisal effect latencies.

### Hierarchical orthographic prediction during natural reading

So far, our results showed that the brain produces distinct prediction error signals at the three levels of linguistic representation, with a temporal order among them. Lastly, as a hallmark of hierarchical prediction, we tested whether the brain utilizes higher-level predictions to inform orthographic predictions. We derived additional oPE variants (Fig. 4A) to test whether such a hierarchical relationship exists. The oPE measure used in the analyses above reflects how strongly the pixels of a given word deviates from a prediction image which is created by averaging word images in a language. The two additional variants of oPE were calculated from prediction images that were the weighted average, rather than simple average, of word images. For context-based oPE, the prediction images were created by weighting each word image by its respective contextual probability from the LLM, assuming that contextual word predictions constrain orthographic predictions. For frequency-based oPE, we weighted each word image by its lexical frequency value, assuming that context-independent lexical predictions constrain orthographic predictions. We compared model fits between models with either context-based, frequency-based, or non-hierarchical oPE. These three models differed only in how oPE was computed, and they were then fitted to participants’ EEG data (downsampled to 250 Hz for computation efficiency). Model fits were cross-validated on a leave-one-block-out basis. For each participant, we first took the median R^2^ of each electrode across folds, and then averaged the median R^2^ across all electrodes. This resulted in three R^2^ values for each of the three models and each participant’s EEG data.

**Fig. 4.**
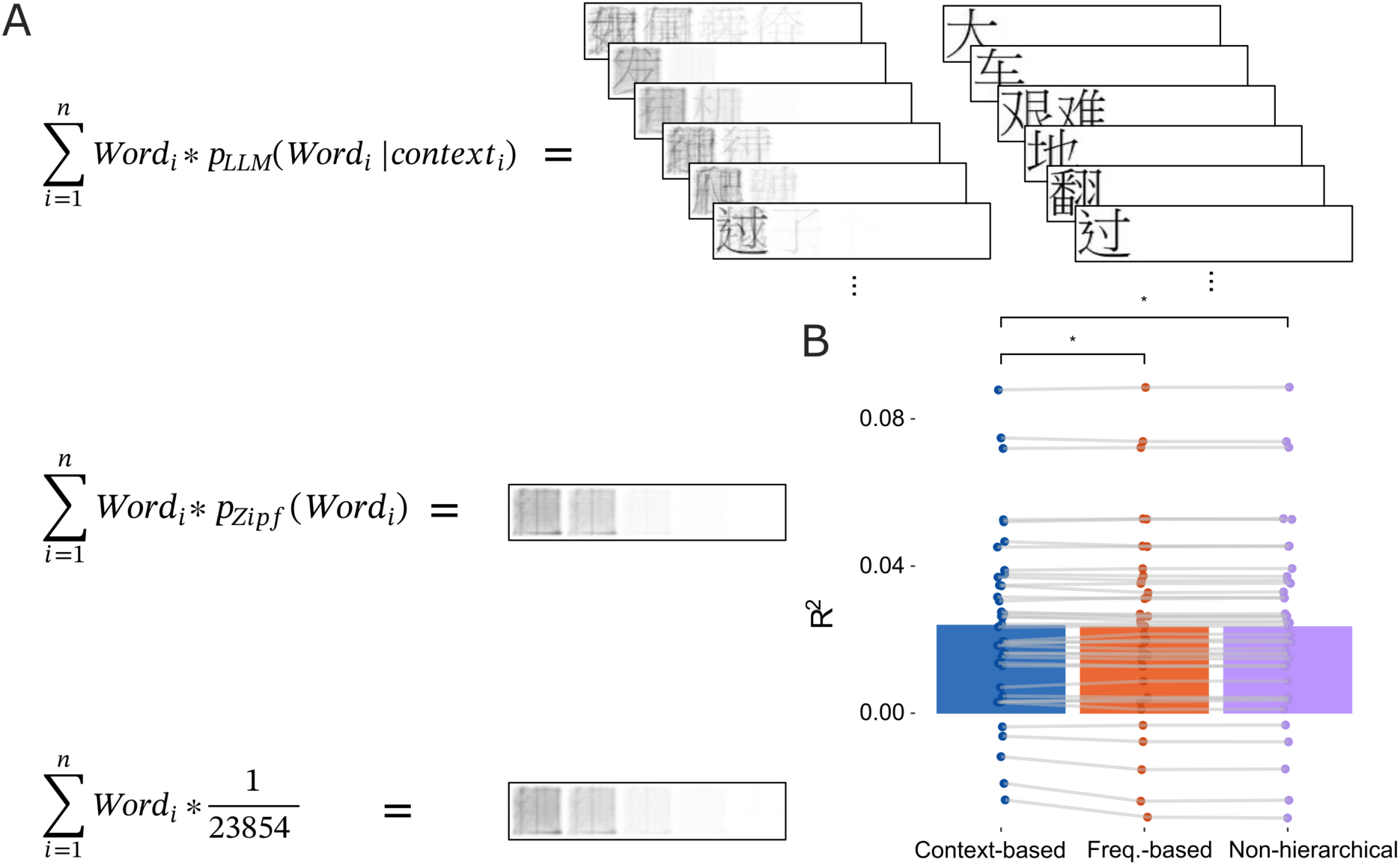
Three types of orthographic prediction images and the result of model-fit comparisons. **(A)** The three variants of oPE were computed based on different prediction images. The top row shows the context-weighted orthographic prediction images on the left, and the respective images of the actual word encountered on the right. oPE computed from these prediction images assume that orthographic predictions are constrained by contextual predictions. Since context changes when new words are encountered, the prediction image also changes dynamically. Note that when a word candidate has high contextual probability compared to others, it becomes more apparent in the resulting orthographic prediction image (e.g., the last prediction image shown). This illustrates the dynamic constraining effect of context on orthographic prediction. The middle row shows the frequency-weighted orthographic prediction image. oPE computed from this image assume that higher-level lexical predictions constrain the lower-level orthographic predictions. The bottom row shows the prediction image derived from simple-average of 23,854 word images. This prediction image was used to derive non-hierarchical oPE. **(B)** Models that differed only in the type of oPE included were compared in terms of their model fit (R^2^) to participants’ EEG data. Each dot shows the cross-validated R^2^ of a participant in the respective model. The same participant is connected by grey lines across the three models. The model with context-based oPE had significantly better fit to participants’ EEG data than the models with frequency-based oPE or non-hierarchical oPE. This finding supports a constraining role of contextual predictions on orthographic predictions.

The models’ cross-validated R^2^ were analyzed with a linear mixed model (Fig. 4B). We found a significant effect of model type, *F*(2,78) = 4.60, *p* = .013. Post-hoc contrasts showed that the model with the context-based oPE (*M* = .0196, *SD* = .0375) explained significantly more variance than the one with the frequency-based oPE (*M* = .019, *SD* = .0386), *t*(78) = 2.7, *p* = .024, and the non-hierarchical model (*M* = .019 *SD* = .0386), *t*(78) = 2.53, *p* = .027. We found no significant difference between the models with frequency-based and non-hierarchical oPE (*p* = .85). The results are consistent with evidence of hierarchical prediction found in listening comprehension (*17*). They also suggest that orthographic predictions are constrained by context-dependent word predictions, thus supporting a hierarchical predictive process in which predictions from higher levels are sent to lower levels during natural reading comprehension.

## Discussion

Taken together, the three types of evidence presented here support the HPP account during natural reading. Each type of evidence also provides specific insights into the neural mechanisms involved during reading. Firstly, we delineated prediction error signals at the orthographic, lexical, and contextual levels by showing that they each had a significant effect on EEG and manifested with distinct topographic distributions. Thus, our findings suggest that the brain generates predictions about upcoming words at multiple levels of linguistic representation, based on their probabilistic features, to support efficient information processing during reading comprehension.

Besides showing that orthographic, lexical, and contextual predictions are implemented, we also show that the three levels of prediction error signals emerged in the expected temporal order. It suggests that the prediction error signals are propagated from the visual-orthographic level to lexical and contextual levels through the predictive hierarchy (*3–5*). Existing models of reading and word recognition often suggest that reading a word activates a cascade of cognitive processes, starting from visual and orthographic information processing to higher-level lexical processes (*44, 45*). Our results reveal a similar cascading process during natural reading. Furthermore, our data suggest relatively rapid lexical access after fixation, as the peak latencies of the word frequency and surprisal effects occured within 300 ms after fixation onset. This is consistent with the proposal that first-order analysis of a word occurs within 300 ms after exposure (*38*), and cognitive processes after this time are elicited by reanalysis and second-order processes (*50*).

In addition to a temporal ordering of signals, we also present evidence that orthographic predictions are constrained by higher-level contextual word predictions. This suggests that higher-level predictions influence lower-level ones and supports the interactive account of the relationship between high-level predictions and low-level visuospatial features during reading (*6*). Moreover, since the surprisal effect’s peak and central latencies are proximate to the average fixation duration of a typical Chinese reader (about 230 ms, *51*), it indicates a rapid update of contextual predictions that subsequently constrain orthographic prediction on the upcoming word. This contextual constraint on low-level predictions have also been shown in listening comprehension (*17*), potentially indicating the same constraining role of contextual predictions on low-level sensory predictions across the two modalities of language comprehension. On the contrary, we did not find evidence indicating that word frequency constrains orthographic predictions. One would expect that higher-level lexical predictions should also constrain lower-level orthographic predictions. This was evident in a recent study in which oPE computed based on the most frequent words fit better to subjects’ EEG data than the "original" oPE in a single-word recognition task (*12*). However, since we used a natural reading task, the availability of rich contextual information could better inform orthographic predictions. Therefore, in the context of natural text reading, readers may select contextual information over lexical frequency for informing orthographic predictions to achieve more efficient processing.

We did not include predictors related to parafoveal word properties, mainly due to the risks of overfitting. However, it is worth noting that the brain may also generate predictions about words located in the parafoveal visual field. For instance, it has been shown that an invalid parafoveal preview of target word resulted in a more negative N400 elicited by pre-target fixation (*35*). Recent evidence also suggested a hierarchical relationship between orthographic and semantic information processing in parafoveal area (*52*). This suggests that the principles of HPP is not specific to foveal processing but may operate across fixations and a wide perceptual span.

Thus, we present evidence from a natural reading paradigm suggesting HPP as the underlying neural mechanism which can facilitate one of the most efficient information-processing skills in human, that is reading. Brain activations are largely consistent with a hierarchical predictive process, indicating multi-level prediction errors that are processed in succession and integrated across levels. This work highlights a hierarchical predictive neural mechanism underlying natural reading, and this mechanism may in fact be fundamental to linguistic processing across modalities.

## Methods

### Participants

Data from 40 participants are reported (35 females; age in years, *M* = 22.9; *SD* = 2.48). Participants were all university students who were skilled readers of simplified Chinese and grew up in Mainland China. They had normal or corrected-to-normal vision, and reported no existing diagnosis of developmental, neurological, and psychiatric disorders. Most participants (n = 39) scored > 50 on the laterality quotient of the Chinese version of Edinburgh Handedness Inventory (*53*, M = 82.9; SD = 18.24), indicating a majority of right-handed participants in the sample. One participant’s data in the first experimental block was not recorded due to human error. The rest of the data from the participant was included in analyses.

The data of four additional participants were excluded from analysis either due to premature termination of experiment (*n* = 1), missing event triggers that compromised synchronization accuracy (*n* = 1), or over 80% of the time windows around fixation onset events being marked as artifacts (*n* = 2, see artifact detection below).

### Stimuli

Experimental stimuli were excerpts from Chinese novels. Half the participants read excerpts from the novel "Mimosa" (*54*) and the other half read "Mr Ma and Son" (*55*). The excerpts contained the first 14,456 and the first 14,614 simplified Chinese characters of each novel, respectively (starting from the beginning), and texts were divided into 400 and 428 sentences, respectively. On a few occasions, the original novels contain English words or Chinese characters that are non-existent in the vocabulary of the Large Language Model (LLM). Those characters and words were replaced with their respective synonyms.

Each character or punctuation mark in the text subtended 1.07° × 1.07° degrees of visual angle. Chinese characters were presented in a Black PMingLiU font (size 40) on a white background. All stimuli were displayed on a Lenovo Legion 24.5 inch LCD monitor (a model G23245FR1 VA panel) refreshing at 144 Hz (1920 x 1080 pixels).

Texts were presented sentence-by-sentence. Sentences typically ended on a full stop, exclamation point, question mark, or ellipsis. In the case where multiple sentences were nested within a pair of quotation marks (e.g., a speech), all materials in the quotation marks were presented in its entirety (i.e., presented as a single sentence). Longer sentences were presented as more than one line on the screen (six lines maximum). An example sentence-layout is shown in Figure S3. The top edge of the first line had a margin of 7.24° from the top edge of the monitor, and each line had a 2.02° margin from the left and 2.5° margin from the right edge of the monitor. In between lines, the bottom edge of the top line was 1.01° apart from the top edge of the bottom line. Each line held a maximum of 32 characters. Characters belonging to the same word were always presented on the same line.

### Procedure

Participants were seated in a sound-attenuated chamber at a viewing distance of 75 cm from the monitor. The experiment was programmed and presented with the Presentation software (*56*). Before the onset of each sentence, participants were asked to maintain fixation on a cross located 1.07° to the left of the first character. Sentence onset was triggered when the eye-tracker detected a fixation within 0.61° from the center of the cross (horizontally and vertically) that lasted 250 ms. Participants read each sentence silently at their own pace and pressed the spacebar once they wished to proceed to the next sentence. This button press would trigger the current sentence to disappear and the left fixation cross to appear again. If sentence onset was not triggered within 10 s of the fixation cross onset, a calibration procedure was prompted to recalibrate the eye-tracker.

The experiment was divided into 20 blocks where each block contained between 15 and 31 sentences. After each block, participants answered two multiple-choice comprehension questions about the content of the preceding block. Each question contained four possible choices with one correct answer. Participants responded by pressing the corresponding key on the keyboard ("D", "F", "J", "K"). Feedback and the correct answer were shown immediately after their response. For example, one of the questions was "Where did Mr. Ma senior first met the priest?". The choices were "D - at his son’s school", "F - at his school", "J - at his brother’s business", and "K - on the ship". An incorrect response would trigger a red background, while a correct response would trigger a green one. The correct answer was shown regardless of the participants’ response. Participants were instructed to answer the questions based on the contents from the previous block only, and to take a break in-between blocks. Before the start of the experiment proper, participants completed a number of practice trials by reading an 8-sentence excerpt from "To Live" (*57*).

### EEG & eye movement recordings

EEG and eye movements were recorded simultaneously during the experiment. EEG was recorded from 128 Ag/AgCl electrodes placed within an ANT Waveguard Original cap with equidistant electrode positions (Advanced Neuro Technology, Enschede, Netherlands). The data was referenced online against an electrode on the vertex (Cz-equivalent) and a clip-on electrode on one earlobe served as ground. A vertical electrooculogram (EOG) electrode was placed under the left eye. Two electrodes located at the lateral orbitofrontal edges of the EEG cap recorded horizontal EOG signals. EEG signals were amplified and recorded with the ANT EEGO HUB amplifier at a sampling rate of 1000 Hz. Electrode impedance was kept below 40 kΩ.

Eye movements were recorded binocularly with a desktop-mounted Eyelink 1000 Plus eye tracker (SR Research) at a sampling rate of 1000 Hz. Participants’ head position was stabilized by a chin rest. A five-point calibration procedure followed by a five-point validation was completed at the beginning of the experiment and whenever a fixation-check failed before the onset of each sentence. Fixations and saccades were detected using Eyelink 1000 tracker’s native algorithm (saccade velocity threshold: 30° degree/s, acceleration threshold: 8000° degree/s^2^).

During the experiment, the stimulus presentation computer sent shared triggers simultaneously to both the EEG and the eye-tracker computer. Offline, both recordings were then synchronized based on these common triggers using the EYE-EEG extension (version 0.99; *21*) for EEGLAB (version 2023.1; *58*). All participants had a synchronization error < 0.41 ms (M = 0.307, SD = 0.045).

### Preprocessing

Preprocessing was done in EEGLAB. EEG data were digitally bandpass filtered using EEGLAB’s Hamming-window sinc FIR filter with cut-off frequencies (−6 dB) at 0.05 and 45.05 Hz (0.1 Hz transition bandwidth). Bad channels, identified by visual inspection, were removed. Independent Component Analysis (ICA) was conducted on a separate copy of the experimental data optimized for ICA training (*19*). Specifically, this copy of the raw dataset was first high- and lowpass-filtered at cutoff frequencies of 3 and 101 Hz, respectively (2 Hz transition bandwidth) and then downsampled to 250 Hz. The same bad channels and intervals that contained large muscle artifacts, were removed. To better capture the brief saccadic spike potential artifact in the ICA training data, EEG data 20 ms before and 10 ms after each saccade (as detected by the eye tracker’s algorithm) was copied and repeatedly appended to the end of the EEG ICA-training data to overweight this artifact for optimal ICA training (*19, 59*). This optimized training dataset was then used for Infomax ICA decomposition, and the resulted ICA weight matrix was applied back to the original dataset. Independent components related to ocular artifacts were then removed by visual inspection. Following ICA, bad channels that were removed before were then interpolated by spherical interpolation and the dataset was re-referenced to the common average reference. Channels that recorded EOG signals were excluded for all further analyses.

### Regression model

For each participant, the continuous EEG data was the dependent variable; and various word properties and control variables constituted a matrix of predictors in a large regression model. Using the unfold toolbox (*20*), the design matrix of the regression model was then time-expanded (*27*), so that the regression coefficients (or “betas”) for each predictor were modelled from 200 ms before and 800 ms after the respective fixation or stimulus event.

To detect bad intervals of the EEG, we then used the "continuousArtifactDetect" function of the ERPLAB toolbox (*60*) as modified for the unfold toolbox. Specifically, a boxcar moving window with a length of 1000 ms was shifted in 500 ms steps across the continuous EEG data. Whenever the window contained a peak-to-peak difference larger than 100 μV, the interval was flagged. Flagged continuous EEG intervals were then excluded from the analysis by setting the corresponding rows of the time-expanded design matrix to 0. For two participants, over 80% of the data around fixation onset events were marked as artifact and these participants were therefore excluded from all analyses. In a final step, the regression model was then solved separately for each EEG channel using the ordinary least-squares method.

The regression model can be described by three separate equations. Each equation denotes a type of event and the corresponding predictors used to model EEG activity around the event. Note that within the model, all predictors are included in the same design matrix and are estimated at the same time:

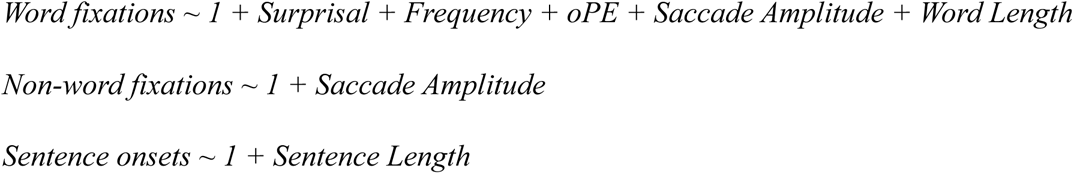

The variables to the left of the tilde denote the event being modelled. "Word fixations" were fixations that landed on a word. Participants made an average of 4419.3 word fixations (*SD* = 1639.6). "Non-word fixations" were fixations that landed on a punctuation mark or on another screen location that did not contain a word. If two eyes were not fixated on the same word, these fixations were also considered as non-word fixations. Participants made an average of 3527.4 non-word fixations (*SD* = 1449.5). "Sentence onsets" were the stimulus onset events when the sentence was first displayed on the screen.

The variables on the right are the predictors used for regressing against EEG at the corresponding event, with the “1” marking the intercept term of the model. Predictors are described in detail below.

#### Surprisal

Word surprisal was estimated by probabilities extracted from the Chinese-Llama 2 Large Language Model (LLM, *22*). This model adopts the LLaMA architecture (*61*), has 13 billion parameters, and was trained with 20 GB of texts from a Chinese corpus. This LLM adopts a tokenizer that was trained with 20,000 Chinese words. Each of these words is represented by a token instead of multiple tokens. Therefore, word probabilities generated by the LLM are probabilities of a whole Chinese word instead of joint probabilities of multiple tokens. This tokenization process was also used as the method of Chinese word segmentation. The LLM and tokenizer were downloaded from Hugging Face’s Transformers package (*62*).

Text from each experimental block was entered to the LLM independently. Hence, the context was reset before every block. The text was first tokenized into word segments (i.e., tokens). The sequence of tokens then passed through the model, and the conditional log-probabilities of each token were extracted from the model. These log-probabilities were then transformed to surprisal by multiplying by −1. Higher surprisal therefore indicates a contextually less probable word. Surprisal values from the two excerpts range from 0.00011 to 17.97 (*M* = 4.16, *SD* =3.06).

#### Frequency

Word frequency was obtained from the SUBTLEX-CH corpus (*24*). We calculated the Zipf value (*63*), a logarithmic measure of word frequency, for each word according to its frequency count and the corpus size. This value transforms the frequency count of word segments to a value proportional to the size of the corpus. It also assigns a value for word segments that have a frequency count of zero in the corpus, assuming that word segments not recorded in SUBTLEX-CH were not completely unseen by the participants. Zipf values were multiplied by −1 so that a higher value represents a less frequent word. Zipf values across the two excerpts range from −7.70 to −1.47 (*M* = −4.87, *SD* = 1.70).

#### Orthographic prediction error (oPE)

We calculated oPE (*13*) with the *EBImage* package (*64*) in the software *R* (*65*). At each word segment position, 55,296 token candidates (vocabulary size of the LLM) were extracted from the LLM. Token candidates that contained non-Chinese characters (e.g. English words, punctuations) were excluded. We also excluded token candidates that were longer than 5 Chinese characters. A total of 23,854 token candidates remained. Each of them was transformed to an image of 265 × 53 pixels in grayscale, with the first character of the word aligned to the left border of the image. Characters were of the same font and size as in the experimental stimuli. In *R*, each image was represented by a 265 × 53 matrix. Numbers in the matrix were continuous values between 1 and 0 that denoted the pixel intensity with 1 = White, 0 = Black, and grey in between. We averaged the matrices of all token candidates to obtain the orthographic prediction image. The difference between the predicted and actual orthographic input was calculated by subtracting the averaged matrix from the matrix of the actual word image. oPE was the sum of all absolute pixel intensities from the resulting difference matrix. Higher oPE value indicates larger difference between the predicted and actual orthographic input. In other words, the oPE calculated by this simple-averaging procedure represents the probability of seeing an orthographic pattern based on all words in the participant’s mental lexicon (i.e., a non-hierarchical oPE). Non-hierarchical oPE range from 925.28 to 2509.33 (*M* = 1113.82, *SD* = 155.41).

Based on the hierarchical predictive processing framework, it is possible that the oPE is influenced by a reader’s prior experience and the preceding context provided by the text. Therefore, for a model comparison analysis, two additional types of oPE were calculated by using weighted-averages, instead of simple-averages, when deriving the orthographic prediction image. The *frequency-weighted* orthographic prediction image was generated by first multiplying each token candidate by their normalized Zipf value, and then summing over all weighted images. oPE calculated from frequency-weighted prediction image (i.e., frequency-based oPE) therefore assumed that orthographic predictions are constrained by the lexical frequency of each word. It therefore reflects a possible hierarchical predictive process where the context-independent probability of each word is used for orthographic prediction.

The *context-dependent* orthographic prediction images were generated by multiplying each token candidate by their normalized context-dependent word probabilities extracted from the LLM (here, a 7-billion parameter version was used for computational efficiency). Since context-dependent word probability changes with context, a unique prediction image was generated for each word segment position. oPE calculated from context dependent orthographic prediction (i.e., context-based oPE) represents a hierarchical predictive process where contextual information constrains orthographic predictions.

#### Saccade amplitude

Saccade amplitude was the size of the saccade preceding the fixation. It has previously been shown that the size of the saccade preceding a given fixation onset is a strong predictor of the FRP waveform, with larger saccades evoking stronger visual brain responses, especially within the early intervals after fixation onset (*27*). This regressor thus captured the variance in FRP that could be attributed to the size of the incoming saccade.

#### Word length

Word length was the number of characters of the fixated word. Longer words tended to have higher oPE values due to more grey pixels in the actual word image. The aim of this regressor was to act as a covariate for the oPE predictor so that the latter better captured variance in EEG attributed to the more subtle effects of orthographic prediction error. We also ran separate control analyses that control for the word length effect on the oPE predictor (see control analyses below). It is important to note that the notion of "word length" here is not equivalent to the number of characters presented in the visual field. This is because the Chinese script does not have explicit word boundaries. In other words, regardless of the length of the currently fixated word, the number of characters present in the visual field at any given time was essentially the same.

#### Sentence length

Sentence length was the number of characters in the sentence. Sentence onset would create an evoked response in EEG which temporally overlaps with the subsequent FRPs (*21*). The amplitude of this evoked response is also expected to depend on the number of characters in the sentence. Hence, sentence onset was modelled by an intercept term and by sentence length to remove the overlapping stimulus-ERP from sentence onset from the FRPs (e.g., *35*).

### Analysis on distinct prediction error signals at each level of hierarchy

From hereafter, *betas* refer to waveforms of beta coefficients from one of the three predictors of interest (i.e., surprisal, frequency, or oPE).

#### Cluster-based permutation with Threshold-Free-Cluster-Enhancement (TFCE)

To establish statistical significance, betas from the word surprisal, frequency, and oPE predictors respectively entered into cluster-based permutation tests (*66*). One-sample t-tests, comparing against 0, were conducted for betas from each time point. The t-values were then enhanced with Threshold-free Cluster Enhancement (TFCE), with parameters E and H set to 2/3 and 2 respectively (*67*). The resulting cluster-enhanced TFCE values then entered into a permutation (*n* = 10,000 random null-hypotheses permutation) test to identify significant clusters.

#### Global Field Power (GFP) and Global Map Dissimilarity (GMD)

Since significant clusters do not establish exact effect latencies (*68*), we also used Global Field Power (GFP) and Global Map Dissimilarity (GMD, *28, 69*) to approximate periods of relatively strong and stable topographic distributions for each predictor’s grand-average betas, thereby providing additional insights into effect latencies.

GFP and GMD are both topographic-level measures (i.e., computed across all electrodes). GFP measures the strength of a topographic map as a function of time. It was equivalent to the standard deviation of betas across the topographic map at a given time point. GMD measures the difference between normalized topographic maps of two consecutive time points. We calculated the squared difference between two consecutive topographic maps divided by their respective GFP. The GMD between two maps is the square root of the averaged squared difference across all electrodes. GMD therefore measures map stability across time regardless of map strength (i.e., GFP). It shows in which time window the predictor produces a stable scalp topography.

We approximated a strong and stable period for each predictor by identifying two GMD minima within a period of high GFP. Although this method is not based on inferential statistics, it establishes an approximate time window of reliable effect based on grand-averaged measures.

#### Control analysis: oPE normalized for word length

The regression model for this control analysis substituted the oPE predictor with oPE normalized for word length ("normalized oPE"). When calculating normalized oPE, only the pixels within the actual word length of the prediction error image were summed together. The summed pixel intensity is then divided by the total number of pixels included in the summation. For example, to compute normalized oPE for a one-character word, pixel intensities within the first character space are summed together. The number is then divided by 2809 which is the total number of pixels that make up a character (53 x 53 pixels).

Normalized oPE ranged from .0024 to .0053 (*M* = .004, *SD* = .00036). Table S1 shows its correlation with other word properties.

#### Control analysis: First-pass fixations on one-character words

The regression model for this control analysis differed from the main analysis on two aspects. Firstly, the "Word fixations" event modelled only first-pass fixations on one-character words. All other fixations were modelled under the "Non-word fixations" event. Secondly, the "Word length" predictor was removed. All other predictors and events modelled remained the same as the main analysis. Table S1 shows the correlations between word properties among all one-character words in the excerpts.

### Analysis of the temporal order of prediction error signals

We identified the peak effect latency with Topographic Component Recognition (*48, 49*). Peak effect latency was established by searching for the maximum correlation between a template map and maps of jackknife-based betas at each time point (i.e., test maps). The template map for each effect was the map at the grand-average GFP maxima. We calculated Pearson correlations between the template map and each of the test maps from 0 ms to 400 ms after fixation onset. This restriction reduced the chance of getting spuriously high correlation coefficient. Importantly, the grand average GFPs of all effects peaked within this range and the same restriction was applied to the three predictors. The peak effect latency of each jackknife-based waveform was defined as when correlation was maximum.

For central effect latency, we first calculated the area and the weighted average position of mass (i.e., moment) between 0 ms and 400 ms with each jackknife-based GFP waveform. We then obtained central effect latency by dividing the moment by the area.

Linear mixed models with predictor type (surprisal, frequency, or oPE) as fixed effect and individual participant as random intercepts were used to test the differences in peak and central effect latencies. All linear mixed models reported were implemented with the *lmerTest* package (*70*) in *R*, fitted with Restricted Maximum Likelihood Estimation, and used the Satterthwaite approximation for degrees of freedom (*71, 72*). Post-hoc comparisons were controlled for familywise error rate with the Bonferroni-Holm method.

### Analysis on hierarchical orthographic prediction during naturalistic reading

We compared the coefficient of determination (R^2^) between three regression models which differed only in how oPE was calculated (see Orthographic prediction error above). All the other regressors were the same as the model used in the main analysis. EEG data were downsampled to 250 Hz before fitted to the three models for computation efficiency. Model comparisons were done based on leave-one-block-out cross-validation which resulted in 19-20 folds per participant (one subject had one block of data missing). We first took the median R^2^ of each electrode across folds. Then, we averaged the R^2^ across all electrodes. Hence, each participant had three cross-validated R^2^ values that estimated the amount of variance explained in the EEG data by each of the three alternative regression models (i.e., non-hierarchical, frequency-based, or context-based oPE). R^2^ was then compared using LMM with model type as a fixed effect and participant as a random intercept.

## Funding

Direct Grant of The Chinese University of Hong Kong 486274300 (UM)

## Authors Contributions

Conceptualization: LC, UM

Methodology: LC, OD, BG, UM

Investigation: LC

Visualization: LC, OD

Funding acquisition: UM, LC

Project administration: UM

Supervision: UM

Writing – original draft: LC

Writing – review & editing: LC, OD, BG, UM

## Notes

### Competing Interest Statement

The authors have declared no competing interest.

