## Supplementary Figures for "Hierarchical Predictive Processing during Natural Reading"

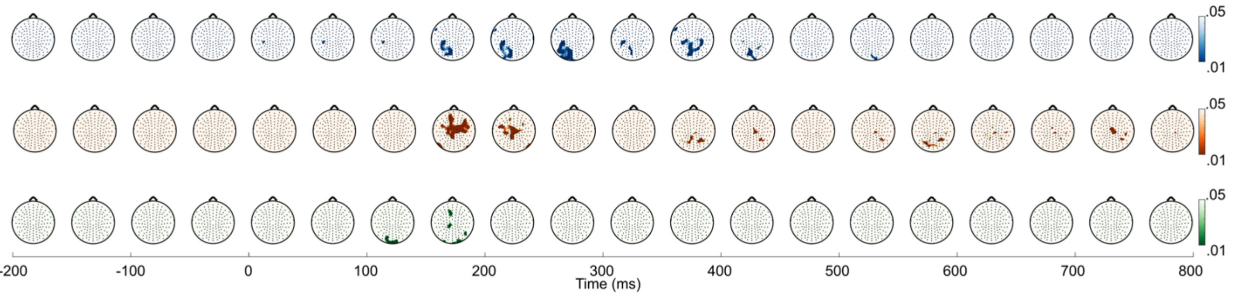


**Fig. S1. P-value maps from cluster-based permutation test of the three predictors.**

All maps have a threshold at $p < .05$. Top row (blue): surprisal effect; Middle row (orange): frequency effect; Bottom row (green): oPE effect.

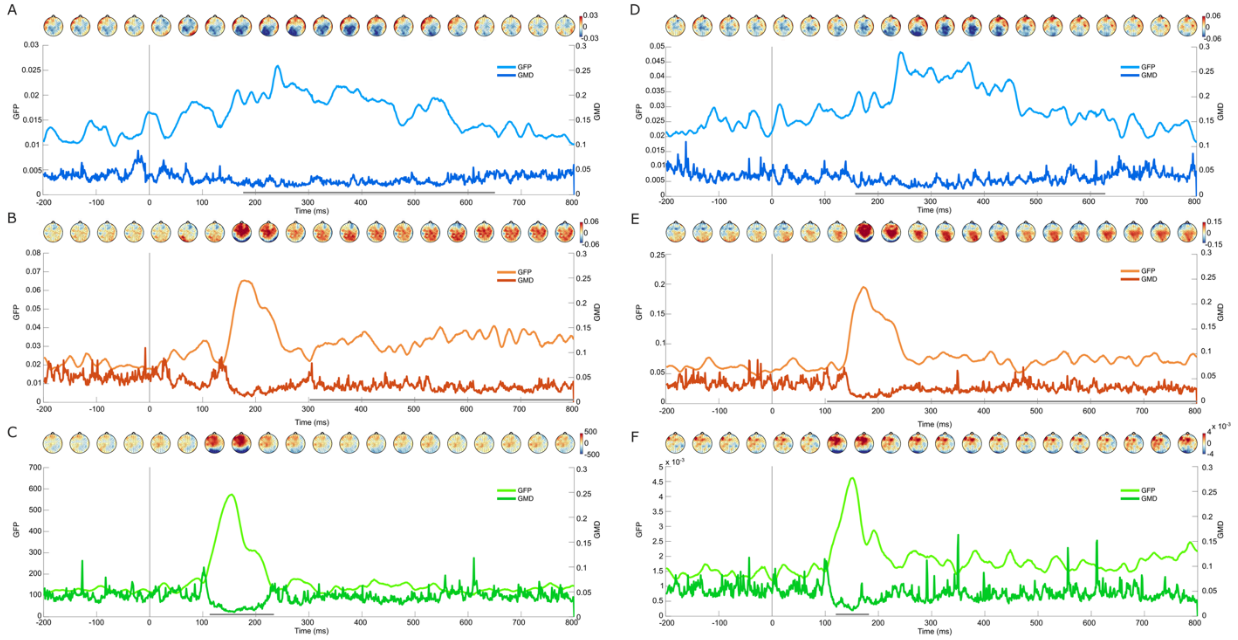

**Fig. S2. Control analyses on the effects of word surprisal, frequency, and oPE.**

**(A, B, C)** Results from the control analysis where normalized oPE was used instead of oPE in the regression models. **(D, E, F)** Results from the control analysis where the three predictors of interest were only used to model first-pass fixation on one-character words. In both control analyses, Global field power (GFP), Global Map Dissimilarity (GMD), and topographic maps show a similar result as in the main analysis. They suggest that the oPE effect found in the main analysis was independent of the word length effect. Top row (blue): surprisal effect; middle row (orange): frequency effect; bottom row (green): oPE effect. Grey horizontal lines indicate significant clusters with $p < .05$.

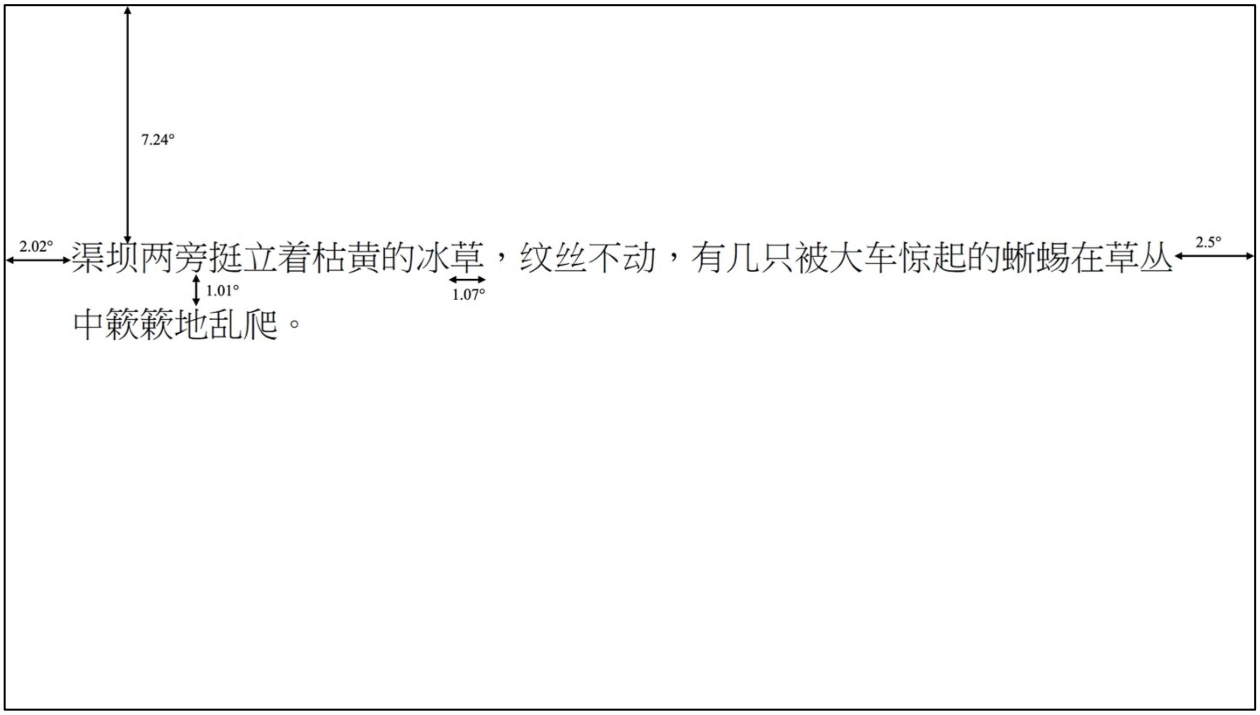


**Fig. S3. Example sentence and sentence layout in the experiment.**

| | Surprisal | Frequency | oPE | Normalized oPE |
|---|---|---|---|---|
| Surprisal | - | - | - | - |
| Frequency | -.26 (-.23) | - | - | - |
| oPE | .24 (-.10) | -.53 (-.30) | - | - |
| Normalized oPE | .09 | .24 | -.27 | - |
| Word length | .25 | -.55 | .90 | -.63 |

**Table S1. Pearson's correlation between word properties in the excerpts.**

Values outside parentheses show the correlations among all words in the excerpts. Values in parentheses show the correlations for all one-character words in the excerpts.

| | All words | | One-character words | |
|---|---|---|---|---|
| | Estimate | *t* | Estimate | *t* |
| Intercept | 200.3 | 28.85*** | 201.0 | 11.49*** |
| Surprisal | 1.5 | 9.47*** | 1.2 | 5.58*** |
| (Inverse) Frequency | 2.3 | 7.26*** | 4.9 | 10.08*** |
| oPE | 0.054 | 9.01*** | 0.05 | 3.21** |
| Saccade amplitude | -1.2 | -12.9*** | -0.43 | -3.44*** |
| Word length | -15.7 | -9.28*** | - | - |

**Table S2. Effects of the fixated word's properties on gaze duration (ms).**

Gaze duration (sum of all first-pass fixations on a word) was modelled with the fixated word's surprisal, inverse lexical frequency, orthographic prediction error (oPE), length, and incoming saccade amplitude as fixed effect. Individual subject and word were modelled as random intercepts. Output on the left modeled gaze durations on words regardless of their length, while output on the right modelled gaze durations only on one-character words. Words that are higher in surprisal, inverse frequency, and oPE received significantly longer gaze durations, corroborating the findings in EEG. *** $p < .001$; ** $p < .01$
